# The SARS-CoV-2 host cell membrane fusion protein TMPRSS2 is a tumor suppressor and its downregulation correlates with increased antitumor immunity and immunotherapy response in lung adenocarcinoma

**DOI:** 10.1101/2021.06.30.450490

**Authors:** Zhixian Liu, Zhilan Zhang, Qiushi Feng, Xiaosheng Wang

**Author notes:** Correspondence to: Xiaosheng Wang.

## Abstract

**Background:** TMPRSS2 is a host cell membrane fusion protein for SARS-CoV-2 invading human host cells. It also has an association with cancer, particularly prostate cancer. However, its association with lung cancer remains insufficiently explored. Thus, an in-depth investigation into the association between TMPRSS2 and lung cancer is significant, considering that lung cancer is the leading cause of cancer death and that the lungs are the primary organ SARS-CoV-2 attacks.

**Methods:** Using five lung adenocarcinoma (LUAD) genomics datasets, we explored associations between *TMPRSS2* expression and immune signatures, cancer-associated pathways, tumor progression phenotypes, and clinical prognosis in LUAD by the bioinformatics approach. Furthermore, we validated the findings from the bioinformatics analysis by performing in vitro experiments with the human LUAD cell line A549 and in vivo experiments with mouse tumor models. We also validated our findings in LUAD patients from Jiangsu Cancer Hospital, China.

**Results:** *TMPRSS2* expression levels were negatively correlated with the enrichment levels of CD8+ T and NK cells and immune cytolytic activity in LUAD, which represent antitumor immune signatures. Meanwhile, *TMPRSS2* expression levels were negatively correlated with the enrichment levels of CD4+ regulatory T cells and myeloid-derived suppressor cells and *PD-L1* expression levels in LUAD, which represent antitumor immunosuppressive signatures. However, *TMPRSS2* expression levels showed a significant positive correlation with the ratios of immune-stimulatory/immune-inhibitory signatures (CD8+ T cells/PD-L1) in LUAD. It indicated that TMPRSS2 levels had a stronger negative correlation with immune-inhibitory signatures than with immune-stimulatory signatures. *TMPRSS2* downregulation correlated with elevated activities of many oncogenic pathways in LUAD, including cell cycle, mismatch repair, p53, and extracellular matrix (ECM) signaling. *TMPRSS2* downregulation correlated with increased proliferation, stemness, genomic instability, tumor advancement, and worse survival in LUAD. In vitro and in vivo experiments validated the association of TMPRSS2 deficiency with increased tumor cell proliferation and invasion and antitumor immunity in LUAD. Moreover, in vivo experiments demonstrated that *TMPRSS2*-knockdown tumors were more sensitive to BMS-1, an inhibitor of PD-1/PD-L1.

**Conclusions:** TMPRSS2 is a tumor suppressor, while its downregulation is a positive biomarker of immunotherapy in LUAD. Our data provide a connection between lung cancer and pneumonia caused by SARS-CoV-2 infection.

## BACKGROUND

The severe acute respiratory syndrome coronavirus 2 (SARS-CoV-2) has infected more than 177 million people and caused more than 3.8 million deaths worldwide as of June 5, 2021 (https://coronavirus.jhu.edu/map.html). SARS-CoV-2 invades host cells using its spike glycoprotein (S) [1], which is composed of S1 and S2 functional domains. S1 binds the angiotensin-converting enzyme 2 (ACE2) for cell attachment, and S2 binds the transmembrane protease serine 2 (TMPRSS2) for membrane fusion [1]. Since TMPRSS2 plays a crucial role in the regulation of SARS-CoV-2 invasion, and cancer patients are susceptible to SARS-CoV-2 infection, an investigation into the role of TMPRSS2 in cancer is significant in the context of the current SARS-CoV-2 pandemic. Previous studies have demonstrated the association between TMPRSS2 and cancer [2-5]. Typically, the TMPRSS2-ERG gene fusion frequently occurs in prostate cancer and is associated with tumor progression [6-8]. In a recent study [3], Katopodis et al. revealed that *TMPRSS2* was overexpressed in various cancers versus their normal tissues. In another study [4], Kong et al. explored *TMPRSS2* expression in lung adenocarcinoma (LUAD) and lung squamous cell carcinoma (LUSC). This study suggested that TMPRSS2 was a tumor suppresser in LUAD for its significant downregulation in LUAD versus normal tissue. A few studies have examined the association between TMPRSS2 and tumor immunity in cancer. For example, Bao et al. [5] investigated *TMPRSS2* expression and its associations with immune and microbiome variates across 33 tumor types. Luo et al. [9] explored the association between *TMPRSS2* expression and immune infiltration in prostate cancer. Despite these prior studies, the associations of TMPRSS2 with tumor immunity, oncogenic signatures or pathways, tumor progression and clinical outcomes in lung cancer remain insufficiently explored.

In this study, we analyzed the associations between *TMPRSS2* expression levels and the enrichment levels of immune signatures in five LUAD cohorts. The immune signatures included CD8+ T cells, NK cells, immune cytolytic activity, CD4+ regulatory T cells, myeloid-derived suppressor cells (MDSCs), and PD-L1. We also analyzed the associations between *TMPRSS2* expression levels and the activities of several oncogenic pathways, including cell cycle, mismatch repair, and p53 signaling. Moreover, we explored the associations between *TMPRSS2* expression and tumor phenotypes (such as proliferation and tumor stemness), genomic features (such as genomic instability and intratumor heterogeneity (ITH)), tumor advancement and prognosis in these LUAD cohorts. Furthermore, we explored the association between *TMPRSS2* expression and the response to cancer immunotherapy. We validated the computational findings by performing in vitro experiments in the human lung cancer cell line A549 and in vivo experiments with mouse tumor models. We also validated our findings in LUAD patients from Jiangsu Cancer Hospital, China. Our study demonstrates that TMPRSS2 is a tumor suppressor while its downregulation can promote antitumor immune response and cancer immunotherapy response. This study may provide insights into the connection between lung cancer and pneumonia caused by SARS-CoV-2 infection.

## METHODS

### Datasets

We downloaded RNA-Seq gene expression profiling (level 3 and RSEM normalized), protein expression profiling, and clinical data for the TCGA-LUAD cohort from the Genomic Data Commons Data Portal (https://portal.gdc.cancer.gov/). We downloaded microarray gene expression profiling (normalized) and clinical data for other four LUAD cohorts (GSE12667 [10], GSE30219 [11], GSE31210 [12], and GSE50081 [13]) from the Gene Expression Omnibus (https://www.ncbi.nlm.nih.gov/geo/). In addition, we collected 100 blood samples from LUAD patients and 20 blood samples from healthy persons from Jiangsu Cancer Hospital, China. According to the diagnosis and treatment guidelines for non-small cell lung cancer (CSCO 2020), LUAD patients in this study were divided into two groups: 50 patients in early stage (stage I) and 50 patients in late stage (stage III-IV). We log2-transformed the RNA-Seq gene expression values before further analyses. A description of these datasets is shown in Supplementary Table S1.

### Gene-set enrichment analysis

We quantified the enrichment levels of immune signatures, pathways, and tumor phenotypes in tumors by the single-sample gene-set enrichment analysis (ssGSEA) [14] of their marker gene sets. The ssGSEA was performed with the R package “GSVA” [14]. The marker gene sets are presented in Supplementary Table S2. We used GSEA [15] to identify KEGG [16] pathways significantly associated with a gene set with a threshold of adjusted *p* value < 0.05. We used WGCNA [17], an R package, to identify gene modules and their associated gene ontology (GO) terms enriched in the high-(upper third) and low-*TMPRSS2*-expression-level (bottom third) LUADs.

### Survival Analysis

We compared overall survival (OS) and disease-free survival (DFS) between the high-(upper third) and low-*TMPRSS2*-expression-level (bottom third) LUAD patients. Kaplan-Meier curves were utilized to display survival time differences, whose significances were evaluated by the log-rank test. We performed the survival analyses using the R package “survival”.

### Statistical analysis

We used the Spearman correlation to evaluate associations between *TMPRSS2* expression levels and ssGSEA scores of gene sets; the Spearman correlation coefficients (ρ) and *p* values were reported. In addition, we used the Pearson correlation to evaluate associations between *TMPRSS2* expression levels and gene or protein expression levels and the ratios of immune signatures; the Pearson correlation coefficients (*r*) were reported. The ratios between immune signatures were the log2-transformed values of the ratios between the geometric mean expression levels of all marker genes in immune signatures. In comparisons of *TMPRSS2* expression levels between two classes of samples, we used the two-tailed Student’s *t* test. We performed the statistical analyses using the R programming software (https://cran.r-project.org/).

### In vitro experiments

#### Antibodies, reagents and cell lines

All antibodies were used at a dilution of 1:1000 unless otherwise specified. Anti-PD-L1 (ab213480), anti-CD8 (ab22378), anti-CD49b (ab181548), anti-MSH6 (ab92471), anti-TMPRSS2 (ab109131) and anti-GAPDH (ab181603) were purchased from Abcam (Burlingame, CA). PE anti-mouse TNF-α antibody (12-7321-81), APC anti-mouse IFN-γ antibody (17-7311-81), APC anti-mouse CD279 (PD-1) antibody (12-9985-81), and APC anti-mouse CD223 (LAG-3) antibody (12-2231-81) were purchased from eBioscience (San Diego, CA). The human lung cancer cell lines A549 were from the American Type Culture Collection. They were cultured in 90% F12K (GIBCO, USA) supplemented with 10% fetal bovine serum in a humidified incubator at 37°C and 5% CO2. NK92 cells (KeyGEN BioTECH, Nanjing, China) were cultured in Alpha MEM (GIBCO, USA) with 2 mM L-glutamine, 1.5 g/L sodium bicarbonate, 0.2 mM inositol, 0.1 mM 2-mercaptoethanol, 0.02 mM folic acid, 100– 200 U/mL recombinant human IL-2 (PeproTech, Rocky Hill, New Jersey, USA), and a final concentration of 12.5% horse serum and 12.5% fetal bovine serum.

#### *TMPRSS2* knockdown with small interfering RNA (siRNA)

A549 cells were transfected with *TMPRSS2* siRNA or control siRNA by using Effectene Transfection Reagent (Qiagen, Hilden, Germany, B00118) according to the manufacturer’s instructions. The medium was replaced after 24 hours incubation with fresh medium, and the cells were maintained for a further 24 hours. Quantitative PCR or Western blotting were used to detect the transfection efficiency. *TMPRSS2* siRNA and control siRNA were synthesized by KeyGEN Biotech (Nanjing, China). Their sequences were as follows: *TMPRSS2* siRNA: 1, 5’-GGAC AUGG GCUA UAAG AAU -3’ (sense) and 5’- AUUC UUAU AGCC CAUG UCC-3’ (antisense); 2, 5’- ACUC CAAG ACCA AGAA CAA -3’ (sense) and 5’- UUGU UCUU GGUC UUGG AGU-3’ (antisense); 3,5’-GGAC UGGA UUUA UCGA CAA-3’(sense) and 5’-UUGU CGAU AAAU CCAG UCC-3’ (antisense); control siRNA: 5’-UUCU CCGA ACGU GUCA CGU dTdT-3’ (sense) and 5’-ACGUGACACGUUCGGAGAAdTdT-3’ (antisense).

#### Lentivirus generation and infection

Lentivirus was prepared according to the manufacturer’s instructions. The heteroduplexes, supplied as 58-nucleotide oligomers, were annealed; the downstream of the U6 promoter was inserted into the pLKO.1 plasmid to generate pLKO.1/ShTMPRSS2. Recombinant and control lentiviruses were produced by transiently transfecting pLKO.1/vector and pLKO.1/ShTMPRSS2, respectively. The lentiviruses were transfected into 293 T cells. After 48 hours, lentiviral particles were collected and concentrated from the supernatant by ultracentrifugation. Effective lentiviral shRNA was screened by infecting these viruses with Lewis cells, and their inhibitory effect on *TMPRSS2* expression was analyzed by quantitative PCR and Western blotting. The lentivirus containing the ShTMPRSS2 RNA target sequences and a control virus were used for the animal study. The coding strand sequence of the shRNA-encoding oligonucleotides was 5’-ACGGGAACGTGACGGTATTTA-3’ for TMPRSS2.

#### Western blotting

A549 cell extracts were lysed by using lysis buffer supplemented with protease inhibitor cocktail immediately before use. Total proteins present in the cell lysates were quantified by using the BCA assay. Proteins were denatured by addition of 6 volumes of SDS sample buffer and boiled at 95°C for 5 min and were then separated by SDS-PAGE. The resolved proteins were transferred onto a nitrocellulose membrane after electrophoresis. The membranes were incubated with 5% skimmed milk in TBS containing 0.1% Tween 20 (TBS-T) for 1 hour to block the non-specific binding and then incubated overnight at 4°C with specific antibodies. After 2 hours incubation with the HRP-labeled secondary antibody, proteins were visualized by enhanced chemiluminescence using a G: BOX chemiXR5 digital imaging system (SYNGENE, UK). The band densities were normalized to the background, and the relative optical density ratios were calculated relative to the housekeeping gene *GAPDH*.

#### Quantitative PCR

The total RNA was isolated by Trizol (Invitrogen, USA) and was reversely transcribed into cDNA using the RevertAid First Strand cDNA Synthesis Kit (Thermo Fisher, USA). Quantitative PCR was performed with the ABI Step one plus Real- Time PCR (RT-PCR) system (ABI, USA) using One Step TB Green(tm) PrimeScript(tm) RT-PCR Kit II (SYBR Green) (RR086B, TaKaRa, JAPAN). Relative copy number was determined by calculating the fold-change difference in the gene of interest relative to GAPTH. The program for amplification was one cycle of 95°C for 5 min, followed by 40 cycles of 95°C for 15 sec, 60°C for 20 sec, and 72°C for 40 sec. The relative amount of each gene was normalized to the amount of *GAPDH*. The primer sequences were as follows: *hTMPRSS2*: 5’-AACT TCAT CCTT CAGG TGTA-3’ (forward) and 5’-TCTC GTTC CAGT CGTCTT-3’ (reverse); *hGAPDH*: 5’- AGAT CATC AGCA ATGC CTCCT-3’ (forward) and 5’-ACAC CATG TATT CCGG GTCAAT-3’ (reverse).

#### Cell proliferation assay

A549 cells were plated in 96-well plates at 3×10^4^ cells per well and maintained in a medium containing 10% FBS. After 24 hours, cell proliferation was determined using the Cell Counting Kit-8 (CCK-8; KeyGEN Biotech, China) following the manufacturer’s instructions. To perform the CCK-8 assay, 10 µl CCK-8 reagent was added to each well and the 96 plates were incubated at 37°C for 2 hours. The optical density was read at 450 nm using a microplate reader. All these experiments were performed in triplicates.

#### Transwell migration and invasion assays

Cell migratory and invasive abilities were assessed using 24 well transwell chambers (Corning, USA) with membrane pore size of 8.0 µm. A549 cells were seeded into the upper chamber without matrigel at 1×10^5^ cells in serum-free medium, while 500 µl medium containing 20% FBS was added to the lower chamber. The chambers were incubated at 37°C and 5% CO_2_ for 24 hours. The cells on the upper chamber were scraped off with cotton-tipped swabs, and cells that had migrated through the membrane were stained with 0.1% crystal violet at 37°C for 30 min. The migrated cells were counted at 200x magnification under the microscope using three randomly selected visual fields. All these experiments were performed in triplicates.

#### Co-culture of tumor cells with NK92 cells

A transwell chamber (Corning, USA) was inserted into a six well plate to construct a co-culture system. A549 cells were seeded on the six well plate at a density of 5×10^4^ cells/well, and NK92 cells were seeded on the membrane (polyethylene terephthalate, pore size of 0.4 µm) of the transwell chamber at a density of 5×10^4^ cells/chamber. Tumor cells and NK92 cells were co-cultured in a humidified incubator at 37°C and 5% CO_2_ atmosphere for 48 hours.

#### EdU proliferation assay

After co-culture of A549 cells with NK92 cells for 48 hours, we measured the proliferation capacity of NK92 cells by an EdU (5- ethynyl-2’-deoxyuridine; Invitrogen, California, USA) proliferation assay. NK92 cells were plated in 96-well plates with a density of 2×10^3^ cells/well with 10 µM EdU at 37°C for 24 hours. The cell nuclei were stained with 4’,6-diamidino-2-phenylindole (DAPI) at a concentration of 1 µg/mL for 20 min. The proportion of NK92 cells incorporating EdU was detected with fluorescence microscopy. All the experiments were performed in triplicates.

### In vivo experiments

#### In vivo mouse models

Lewis tumor cells were transduced with ShCon (scramble) or ShTMPRSS2 lentivirus and selected by puromycin for 7 days. The stably transfected Lewis tumor cells (1×107/ml) were subcutaneously injected into the right armpit of recipient mice after shaving the injection site. After 5 days, when the tumor volume was approximately 4-5 mm3, the mice were randomly divided into six groups, with half of the ShCon and ShTMPRSS2 mice treated with 150 U/L PD1/PDL1 inhibitor BMS-1 (concentration 500 mg/mL; i.p.) (MCE Cat. No. HY-19991) every 3 days. The tumors were isolated from mice after 15 days. Tumor volumes did not exceed the maximum allowable size according to the LJI IACUC animal experimental protocol. The tumor volume was measured every 3 days after the tumor appeared on the fifth day and was calculated as follows: V = 1/2 × width2 × length.

#### Isolation of tumor-infiltrating lymphocytes (TILs)

After the tumor tissues were separated aseptically and rinsed with cold PBS for 3 times, they were excised and chopped with tweezers and scissors and were then digested with 2 mg/mL collagenase (type IV, sigma V900893) for 45 min, until no tissue mass was visible. Following digestion, lymphocytes were separated with lymphocyte separation medium, washed with PBS, and counted. The specific protocol was as follows: tumors were filtered through 70 µM cell strainers, and the cell suspension was washed twice in culture medium by centrifugation at 1500 rpm and 4°C for 10 min. After the washing, the cells were resuspended with PBS and were layered over 3 mL of 30%-100% gradient percoll (Beijing Solarbio Science & Technology, Beijing, China); this was followed by centrifugation at 2600 rpm for 25 min at 25°C. The enriched TILs were obtained at the interface as a thin buffy layer, were washed with PBS three times, and finally were resuspended in FACS staining buffer for further staining procedures.

#### Flow cytometry

TILs were stained with CD8 (eBioscience, 11-0081-81), CD49b (eBioscience, 11-5971-81), PD-1 (eBioscience, 12-9985-81), and LAG3 (eBioscience, 12-2231-81) and were analyzed by flow cytometry. TILs were restimulated with cell stimulation cocktail (eBioscience, San Diego, California, USA), and the expression of IFN-γ and TNF-α (Biolegend) was analyzed by flow cytometry. Staining for cell surface markers was performed by incubating cells with antibody (1:100 dilution) in FACS buffer (0.1% BSA in PBS) for 30 min at 4°C. Surface markers of intracellular cytokines (IFN-γ (eBioscience, 17-7311-81) and TNF-α (eBioscience, 12-7321-81)) were stained before fixation/permeabi-lization (Intracellular Fixation & Permeabilization Buffer Set, ThermoFisher).

#### Immunofluorescence of CD8, CD49b and PD-L1

Paraffin-embedded mice tumor tissue sections (3 µm thick) were subjected to immunofluorescence with CD8 (Abcam, ab22378), CD49b (Abcam, ab181548), or PD-L1 (Abcam, ab2134808) primary antibodies. Before immunostaining, tumor tissue sections were deparaffinized with xylene, rehydrated and unmasked in sodium citrate buffer (10 mM, pH 6.0), and treated with a glycine solution (2 mg/mL) to quench autofluorescence. After antigen retrieval, 3% H2O2-methanol solution blocking inactivated enzymes, and goat serum blocking, tissue slides were incubated in wet box for 2 hours at 37°C with anti-CD8, CD49b, or anti-PD-L1 rabbit primary antibodies (1:100 dilution) in blocking solution, and were then dropped with FITC (1:100 dilution) secondary antibody 50-100ul and incubated at 37° for 1 hour in the dark. The immunolabeled slides were examined with a fluorescence microscope after nuclear counterstaining with DAPI. Green, red and blue channel fluorescence images were acquired with a Leica DFC310 FX 1.4-megapixel digital color camera equipped with LAS V.3.8 software (Leica Microsystems, Wetzlar, Germany). Overlay images were reconstructed by using the free-share ImageJ software.

## RESULTS

### Associations between *TMPRSS2* expression and immune signatures in LUAD

We found that *TMPRSS2* had a significant negative expression correlation with the infiltration levels of CD8+ T cells, which represent the adaptive antitumor immune response, in three of the five LUAD cohorts (Spearman correlation, *p* < 0.05) (Figure 1A). *TMPRSS2* expression levels were also significantly and negatively correlated with the infiltration levels of NK cells, which represent the innate antitumor immune response, in two LUAD cohorts (*p* < 0.05) (Figure 1A). Moreover, *TMPRSS2* expression levels were negatively correlated with immune cytolytic activity, a marker for underlying immunity [18], in all the five LUAD cohorts. Meanwhile, *TMPRSS2* had a significant negative expression correlation with *PD-L1* in the five LUAD cohorts (Figure 1A). *TMPRSS2* expression levels were negatively correlated with the infiltration levels of CD4+ regulatory T cells and MDSCs in four LUAD cohorts, which represent tumor immunosuppressive signatures (Figure 1A). Taken together, these results suggest a significant negative association between TMPRSS2 abundance and immune infiltration levels in LUAD. Interestingly, *TMPRSS2* expression levels showed a significant positive correlation with the ratios of immune-stimulatory/immune-inhibitory signatures (CD8+ T cells/PD-L1) consistently in the five LUAD cohorts (Pearson correlation, *p* < 0.05) (Figure 1B). It indicated that TMPRSS2 levels had a stronger negative correlation with immune-inhibitory signatures than with immune-stimulatory signatures. Furthermore, we found that the ratios of immune-stimulatory/immune-inhibitory signatures were positively correlated with DFS in the TCGA-LUAD cohort (log-rank test, *p* = 0.01) (Figure 1C).

**Figure 1.**
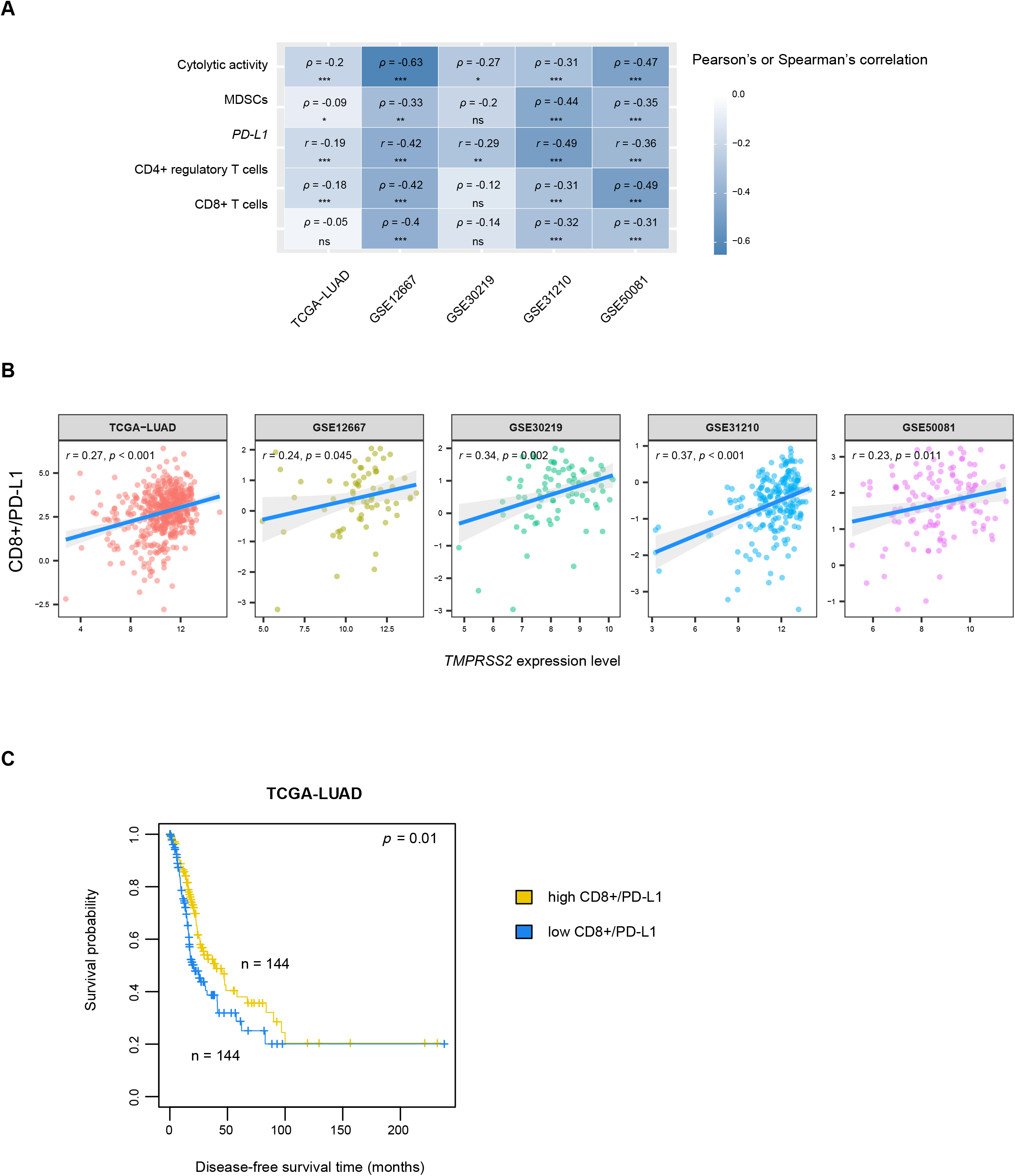
Association between *TMPRSS2* expression and immune signatures in LUAD. **(A)** Correlations between *TMPRSS2* expression levels and the enrichment levels of CD8+ T cells and NK cells, immune cytolytic activity, *PD-L1* expression levels, and the enrichment levels of CD4+ regulatory T cells and MDSCs in five LUAD cohorts. The Spearman or Pearson correlation coefficients (ρ or *r*) and *p* values are shown. **(B)** Pearson correlations between *TMPRSS2* expression levels and the ratios of immune-stimulatory/immune-inhibitory signatures (CD8+/PD-L1) in LUAD. **(C)** Kaplan-Meier survival curves showing a better disease-free survival in LUAD patients with high ratios of CD8+/PD-L1 (upper third) than those with low ratios of CD8+/PD-L1 (bottom third). The log-rank test *p* value is shown. * *p* < 0.05, ** *p* < 0.01, *** *p* < 0.001, ^ns^ *p* ≥ 0.05. They also apply to the following figures.

### Associations between *TMPRSS2* expression and oncogenic pathways, tumor phenotypes and prognosis in LUAD

We found that *TMPRSS2* expression levels were inversely correlated with the activities of the cell cycle, mismatch repair, and p53 signaling pathways in the five LUAD cohorts (Spearman correlation, *p* < 0.001) (Figure 2A). Moreover, *TMPRSS2* showed a negative expression correlation with *MKI67*, a tumor proliferation marker, in the five LUAD cohorts (Pearson correlation, *p* < 0.001) (Figure 2B). Tumor stemness indicates a stem cell-like tumor phenotype representing an unfavorable prognosis in cancer [19]. We observed that *TMPRSS2* expression levels were inversely correlated with tumor stemness scores in these LUAD cohorts (Spearman correlation, *p* < 0.001) (Figure 2C).

**Figure 2.**
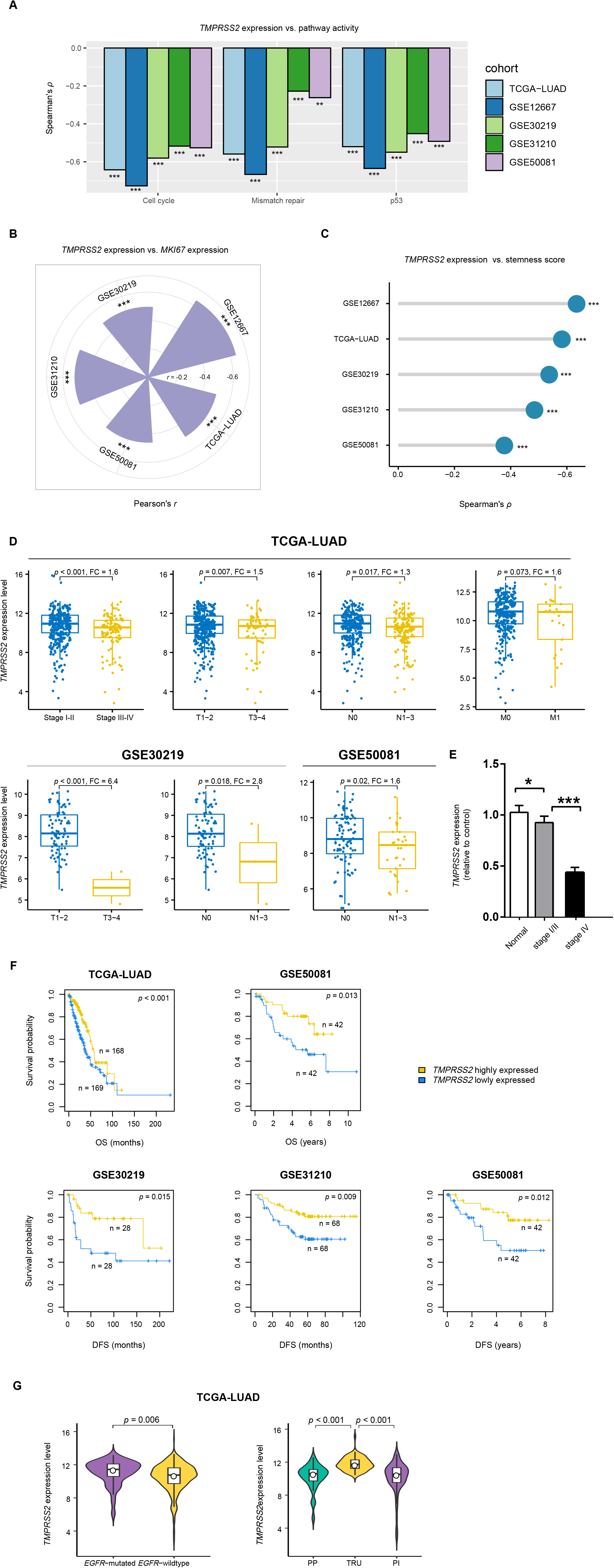
Associations between *TMPRSS2* expression and oncogenic pathways, tumor phenotypes and prognosis in LUAD. The inverse correlations between *TMPRSS2* expression levels and the activities of oncogenic pathways **(A)**, *MKI67* expression levels **(B)**, and stemness scores **(C)** in LUAD. The Spearman or Pearson correlation coefficients (ρ or *r*) and *p* values are shown. **(D)** Comparisons of *TMPRSS2* expression levels between late-stage (Stage III-IV) and early-stage (Stage I-II), between large-size (T3-4) and small-size (T1-2), and between N1-3 (lymph nodes) and N0 (without regional lymph nodes) LUADs. The Student’s *t* test *p* values and fold change (FC) of mean *TMPRSS2* expression levels are shown. **(E)** The lung cancer data from Jiangsu Cancer Hospital showing that *TMPRSS2* expression levels are significantly lower in late-stage (Stage IV) than in early-stage (Stage I-II) LUADs. **(F)** Kaplan-Meier survival curves showing that low-*TMPRSS2*-expression-level (bottom third) LUAD patients have worse OS and/or DFS than high-*TMPRSS2*-expression-level (upper third) LUAD patients. The log-rank test *p* values are shown. OS, overall survival. DFS, disease-free survival. **(G)** Comparisons of *TMPRSS2* expression levels between *EGFR*-mutated and *EGFR*-wildtype LUADs and between three LUAD transcriptional subtypes. TRU, terminal respiratory unit. PI, proximal-inflammatory. PP, proximal-proliferative.

We detected that *TMPRSS2* expression levels significantly decreased with tumor advancement in LUAD (Figure 2D). For example, in the TCGA-LUAD cohort, *TMPRSS2* expression levels were significantly lower in late-stage (Stage III-IV) than in early-stage (Stage I-II) LUADs (Student’s *t* test, *p* < 0.001; fold change (FC) = 1.6), in large-size (T3-4) than in small-size (T1-2) LUADs (*p* = 0.007; FC = 1.5), in LUADs with lymph nodes (N1-3) than in those without regional lymph nodes (N0) (*p* = 0.02; FC = 1.3), and in LUADs with metastasis (M1) than in those without metastasis (M0) (*p* = 0.07; FC = 1.6). In other two LUAD cohorts (GSE30219 and GSE50081) with tumor size and lymph nodes data available, *TMPRSS2* expression levels were also significantly lower in large-size than in small-size LUADs (*p* < 0.001; FC = 6.4) in GSE30219 and were significantly lower in N1-3 than in N0 LUADs in both GSE30219 (*p* = 0.02; FC = 2.83) and GSE50081 (*p* = 0.02; FC = 1.6) (Figure 2D). Furthermore, the lung cancer data from Jiangsu Cancer Hospital supported that *TMPRSS2* expression levels were reduced in late-stage (Stage IV) than in early-stage (Stage I-II) LUADs (*p* < 0.001; FC = 1.6) (Figure 2E). Survival analyses showed that *TMPRSS2* downregulation was correlated with worse OS and/or DFS in these LUAD cohorts (log-rank test, *p* < 0.05) (Figure 2F).

It has been shown that *EGFR-*mutated LUADs have a better prognosis than *EGFR*-wildtype LUADs [20]. We found that *TMPRSS2* was more lowly expressed in *EGFR*-wildtype than in *EGFR*-mutated LUADs (*p* = 0.006; FC = 1.5) (Figure 2G). Besides, LUAD harbors three transcriptional subtypes: terminal respiratory unit (TRU), proximal-inflammatory (PI), and proximal-proliferative (PP), of which TRU has the best prognosis [21]. We found that *TMPRSS2* expression levels were the highest in TRU (TRU versus PP: *p* = 8.68 × 10^−14^, FC = 2.98; TRU versus PI: *p* = 1.07 × 10^−11^, FC = 3.16) (Figure 2G).

Taken together, these results suggest that TMPRSS2 downregulation is associated with worse outcomes in LUAD.

### Association between *TMPRSS2* expression and genomic instability in LUAD

Genomic instability plays prominent roles in cancer initiation, progression, and immune invasion [22] by increasing TMB [23] and aneuploidy or somatic copy number alterations [24]. In the TCGA-LUAD cohort, *TMPRSS2* expression levels had a negative correlation with TMB (Spearman correlation, ρ = -0.31; *p* = 2.58 × 10^−12^) (Figure 3A). Homologous recombination deficiency (HRD) may promote chromosomal instability and aneuploidy levels in cancer [25]. We found that *TMPRSS2* expression levels were inversely correlated with HRD scores [25] in LUAD (ρ = -0.27; *p* = 5.76 × 10^−10^) (Figure 3B). DNA damage repair (DDR) deficiency can lead to genomic instability [26]. Knijnenburg et al. [25] identified deleterious gene mutations for nine DDR pathways in TCGA cancers. We divided LUAD into pathway-wildtype and pathway-mutated subtypes for each of the nine DDR pathways. The pathway-wildtype indicates no deleterious mutations in any pathway genes, and the pathway-mutated indicates at least a deleterious mutation in pathway genes. Interestingly, we found that *TMPRSS2* expression levels were significantly lower in the pathway-mutated subtype than in the pathway-wildtype subtype for seven DDR pathways (*p <* 0.05; FC > 1.5) (Figure 3C). The seven pathways included base excision repair, Fanconi anemia, homologous recombination, mismatch repair, nucleotide excision repair, translesion DNA synthesis, and damage sensor. These results suggest a correlation between TMPRSS2 downregulation and DDR deficiency.

**Figure 3.**
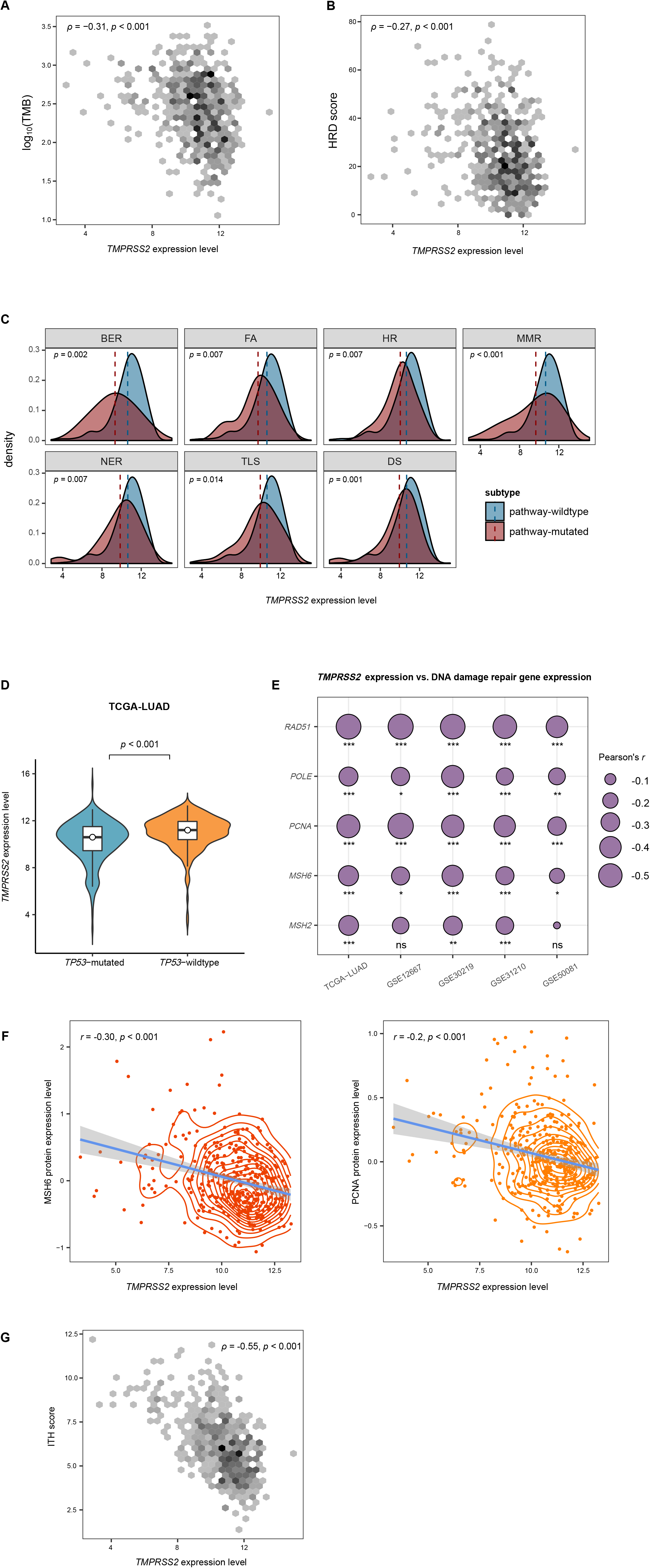
Association between *TMPRSS2* expression and genomic instability in LUAD. Spearman correlations between *TMPRSS2* expression levels and tumor mutation burden (TMB) **(A)** and homologous recombination deficiency (HRD) scores **(B)** in TCGA-LUAD. TMB is the total somatic mutation count in the tumor. The HRD scores were obtained from the publication [25]. **(C)** Comparisons of *TMPRSS2* expression levels between pathway-wildtype and pathway-mutated LUAD subtypes for seven DNA damage repair (DDR) pathways in TCGA-LUAD. The pathway-wildtype indicates no deleterious mutations in any pathway genes, and the pathway-mutated indicates at least a deleterious mutation in pathway genes. BER, base excision repair. FA, Fanconi anemia. HR, homologous recombination. MMR, mismatch repair. NER, nucleotide excision repair. TLS, translesion DNA synthesis. DS, damage sensor. **(D)** Comparisons of *TMPRSS2* expression levels between *TP53*-mutated and *TP53*-wildtype LUADs. Expression correlations between *TMPRSS2* and DDR-associated genes **(E)** and proteins **(F)** in LUAD. **(G)** Spearman correlation between *TMPRSS2* expression levels and intratumor heterogeneity (ITH) scores. The ITH scores were evaluated by the DEPTH algorithm [29].

*TP53* mutations often leads to genomic instability because of the important role of p53 in maintaining genomic stability [27]. We found that *TMPRSS2* displayed significantly lower expression levels in *TP53*-mutated than in *TP53*-wildtype LUADs (*p* = 0.006; FC = 1.5) (Figure 3D). Moreover, we found numerous DDR-associated genes having significant negative expression correlations with *TMPRSS2* in these LUAD cohorts (Pearson correlation, *p* < 0.05), including *MSH2, MSH6, POLE, PCNA*, and *RAD51* (Figure 3E). Furthermore, we observed significant negative expression correlations between *TMPRSS2* and DNA mismatch repair proteins MSH6 (Pearson correlation, *r* = -0.30; *p* = 6.6 × 10^−9^) and PCNA (*r* = -0.25; *p* = 1.5 × 10^−6^) in the TCGA-LUAD cohort (Figure 3F). These results indicated an association between TMPRSS2 downregulation and the upregulation of DDR molecules, the signature of increased genomic instability.

Genomic instability can promote tumor heterogeneity, which is associated with tumor progression, immune evasion, and drug resistance [28]. We used the DEPTH algorithm [29] to score ITH for each TCGA-LUAD sample and found a significant negative correlation between *TMPRSS2* expression levels and ITH scores in LUAD (ρ = -0.55; *p* < 0.001) (Figure 3G). It indicates a significant association between TMPRSS2 downregulation and increased ITH in LUAD.

Taken together, these results suggest that TMPRSS2 downregulation is associated with enhanced genomic instability in LUAD.

### Co-expression networks of *TMPRSS2* in LUAD

We found 150 and 135 genes having strong positive and negative expression correlations with *TMPRSS2* in the TCGA-LUAD cohort, respectively (Pearson correlation, |*r*| > 0.5) (Figure 4A; Supplementary Table S3). GSEA [14] revealed that the cell cycle, p53 signaling, mismatch repair, and homologous recombination pathways were significantly associated with the 135 genes with strong negative expression correlations with *TMPRSS2*. This conforms to the previous findings that *TMPRSS2* downregulation was correlated with increased activities of these pathways.

**Figure 4.**
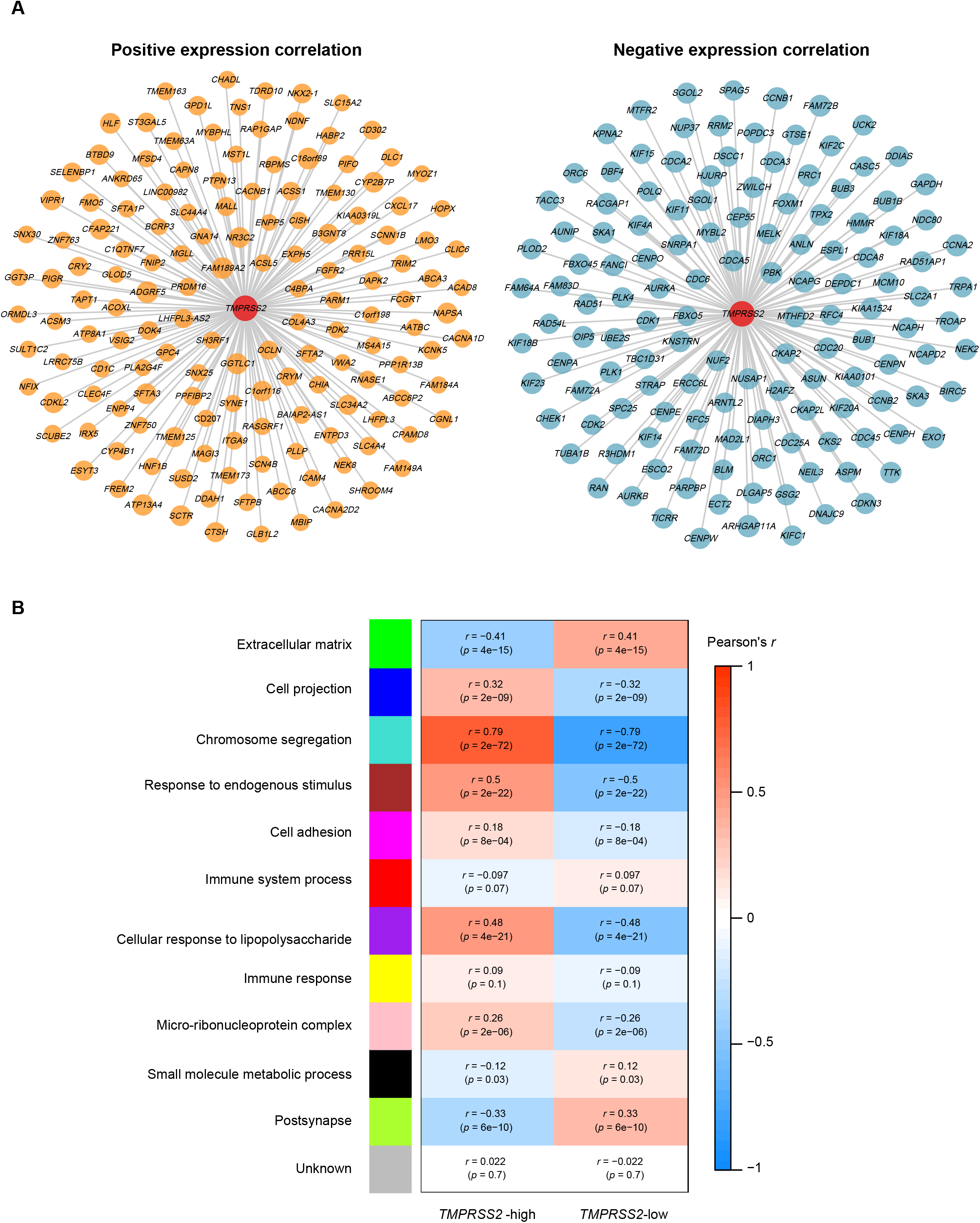
Co-expression networks of *TMPRSS2* in LUAD. **(A)** 150 and 135 genes having strong positive and negative expression correlations with *TMPRSS2* in TCGA-LUAD, respectively (|*r*| > 0.5). **(B)** Gene modules and their representative gene ontology terms highly enriched in high-(upper third) and low-*TMPRSS2*-expression-level (bottom third) LUADs identified by WGCNA [17].

WGCNA [17] identified six gene modules (indicated in blue, turquoise, brown, magenta, purple, and pink color, respectively) highly enriched in the high-*TMPRSS2*-expression-level LUADs. The representative GO terms associated with these modules included cell projection, chromosome segregation, response to endogenous stimulus, cell adhesion, cellular response to lipopolysaccharide, and micro-ribonucleoprotein complex. In contrast, three gene modules (indicated in green, black, and green-yellow color, respectively) were highly enriched in the low-*TMPRSS2*-expression-level LUADs (Figure 4B). The representative GO terms for these modules included extracellular matrix (ECM), small molecule metabolic process, and postsynapse (Figure 4B). The ECM signature plays a crucial role in driving cancer progression [30]. Its upregulation in the low-*TMPRSS2*-expression-level LUADs is in accordance with the correlation between *TMPRSS2* downregulation and LUAD progression.

### Experimental validation of the bioinformatics findings

To validate the findings from the bioinformatics analysis, we performed in vitro experiments with the human LUAD cell line A549 and in vivo experiments with mouse tumor models. We found that *TMPRSS2* knockdown markedly promoted proliferation and invasion potential in A549 cells (Figure 5A) and increased tumor volume and progression in Lewis tumor mouse models (Figure 5B). This is consistent with the previous results showing that TMPRSS2 downregulation is associated with tumor progression and unfavorable prognosis in LUAD. Furthermore, in vitro experiments showed that MSH6 expression was upregulated in *TMPRSS2*-knockdown versus *TMPRSS2*-wildtype A549 cells (Figure 5C). This is in agreement with the previous finding of the significant negative correlation between *TMPRSS2* expression levels and MSH6 abundance in LUAD.

**Figure 5.**
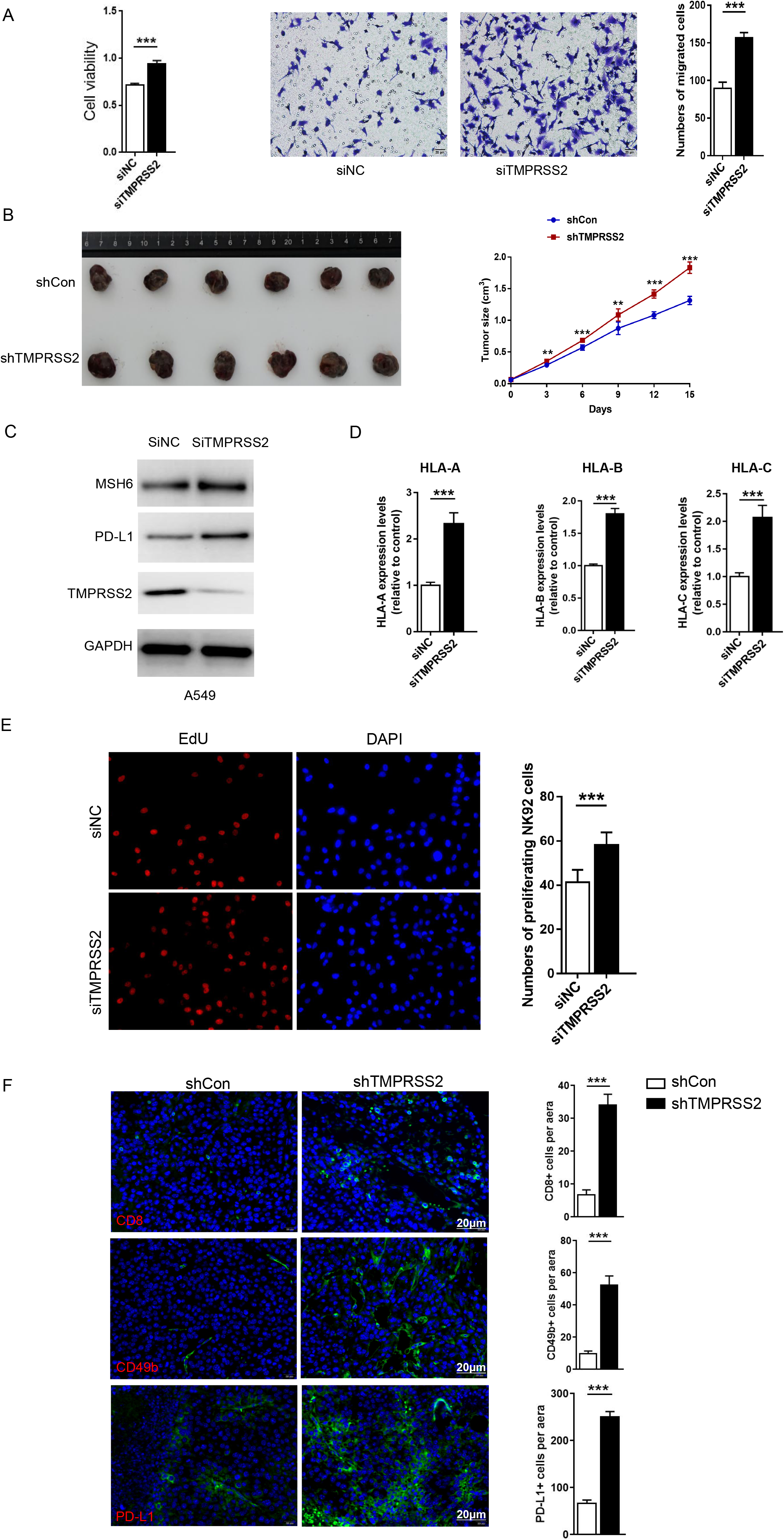
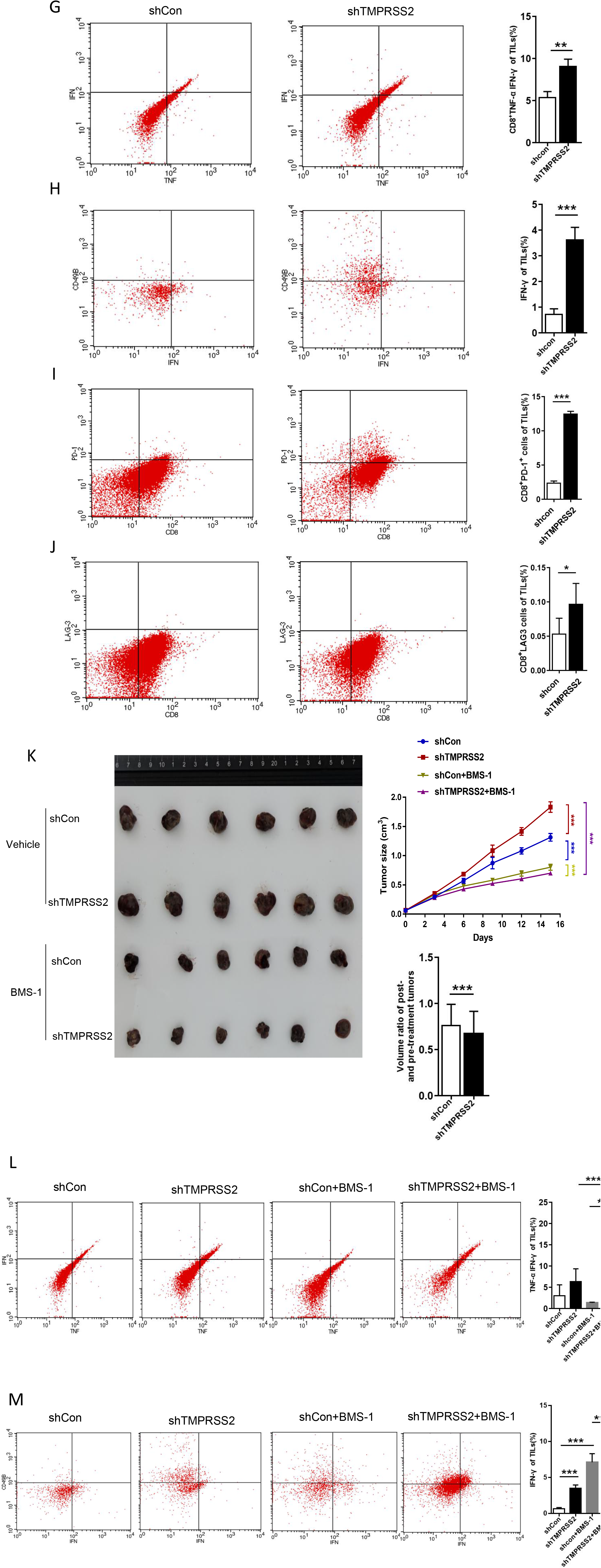
In vivo and in vitro experimental validation of the bioinformatics findings. TMPRSS2-knockdown tumors display increased tumor-infiltrating lymphocytes, expression of immune checkpoint molecules, and sensitization to immune checkpoint inhibitors. **(A)** *TMPRSS2* knockdown markedly promoted proliferative and invasive abilities of A549 cells. **(B)** *TMPRSS2* knockdown increased tumor volume and progression in Lewis tumor mouse models. Lewis tumor cells transfected with ShCon or ShTMPRSS2 lentivirus were subcutaneously injected into mice. The tumor volumes were measured every three days from the fifth day to the fifteenth. Data represent mean□±□SEM. SEM, standard error of mean. ShTMPRSS2 versus ShCon group, *n* = 6 for each group, two-tailed Student’s *t* test, * *p* < 0.05, ** *p* < 0.01, *** *p* < 0.001. **(C)** *TMPRSS2* knockdown increased MSH6 expression in A549 cells, as evidenced by Western blotting. **(D)** *TMPRSS2* knockdown enhanced the expression of MHC class I genes (*HLA-A, HLA-B*, and *HLA-C*) in A549 cells, as evidenced by real-time qPCR. **(E)** NK cells co-cultured with *TMPRSS2*-knockdown A549 cells showing higher proliferation capacity than NK cells co-cultured with *TMPRSS2*-wildtype A549 cells, as evidenced by the EDU proliferation assay. **(F)** CD8, CD49b, and PD-L1 immunofluorescence staining in Lewis orthotopic tumors and H-score analysis. ShTMPRSS2 versus shCon group, *n* = 6 for each group, two-tailed Student’s *t* test, *** *p* < 0.001. **(G-J)** Comparisons of TNF-α, IFN-γ, PD-1, and LAG3 expression on CD8+ T cells from tumor-infiltrating lymphocytes (TILs) in tumor-bearing mice between *TMPRSS2*-knockdown and *TMPRSS2*-wildtype group (ShTMPRSS2 versus ShCon group, *n* = 6 for each group, two-tailed Student’s *t* test, * *p* < 0.05, ** *p* < 0.01, *** *p* < 0.001). TILs were stained with CD3, CD8, TNF-α, and IFN-γ and were then analyzed by flow cytometry. Lymphocytes were gated according to forward scatter and side scatter. CD3 and CD8 staining was used to identify CD8+ T cells. **(K-M)** *TMPRSS2*-knockdown tumors formed by subcutaneous injection of Lewis cells, as mentioned in **(B)**. shCon and shTMPRSS2 tumor-bearing mice were divided into vehicle and BMS-1 groups. The vehicle and BMS-1 groups of mice were treated with solvent and BMS-1, respectively.**(K)** Representative images of tumor-bearing mice shown on the left. The right graph showing the change of tumor size in the tumor-bearing mice over time. Data represent mean□±□SEM (*n* = 6 for each group, two-tailed Student’s *t* test, * *p* < 0.05, ** *p* < 0.01, *** *p* < 0.001); Comparison of the volume ratios of mice tumors after and before treatment with BMS-1 between *TMPRSS2*-knockdown and *TMPRSS2*-wildtype groups (two-tailed Student’s *t* test, *** *p* < 0.001). Comparisons of TNF-α **(L)** and IFN-γ **(M)** expression on CD8+ T cells from TILs in tumor-bearing mice (*n* = 6 for each group, two-tailed Student’s *t* test, * *p* < 0.05, ** *p* < 0.01, *** *p* < 0.001).

Our bioinformatics analysis revealed a significant inverse correlation between TMPRSS2 abundance and immune infiltration levels in LUAD. Consistently, the MHC class I genes (*HLA-A, HLA-B*, and *HLA-C*) showed significantly higher expression levels in *TMPRSS2*-knockdown than in *TMPRSS2*-wildtype A549 cells, demonstrated by real-time qPCR (Figure 5D). NK cells co-cultured with *TMPRSS2*-knockdown A549 cells displayed significantly stronger proliferation ability than NK cells co-cultured with *TMPRSS2*-wildtype A549 cells, evident by the EdU proliferation assay (Figure 5E). Furthermore, in vivo experiments showed that infiltration of CD8+ T cells and NK cells significantly increased in *TMPRSS2*-knockdown tumors (Figure 5F). Moreover, on CD8+ T cells from TILs in *TMPRSS2*-knockdown tumors, the expression of TNF-α and IFN-γ were significantly upregulated (Figure 5G, H), indicating that *TMPRSS2* knockdown can enhance the activity of CD8+ TILs. Meanwhile, the expression of PD-1 and LAG3 also significantly increased on CD8+ TILs in *TMPRSS2*-knockdown tumors (Figure 5I, J), indicating that *TMPRSS2* deficiency can also promote the exhaustion of CD8+ TILs.

Our bioinformatics analysis revealed a significant negative correlation between *TMPRSS2* and *PD-L1* expression levels. This result was confirmed by both in vitro and in vivo experiments; knockdown of *TMPRSS2* increased PD-L1 expression in A549 cells, as evidenced by Western blotting (Figure 5C); *TMPRSS2*-knockdown tumors had significantly enhanced PD-L1 expression (Figure 5F). Furthermore, bioinformatics analysis revealed a significant positive correlation between *TMPRSS2* expression levels and the ratios of CD8+ T cells/PD-L1. This was confirmed by that *TMPRSS2*-knockdown tumors displayed a higher level of increases in CD8+ T cell infiltration than in PD-L1 abundance (Figure 5F). Because PD-L1 expression is a predictive biomarker of response to immune checkpoint inhibitors (ICIs) in cancer [31], we anticipated that knockdown of *TMPRSS2* would promote the response to ICIs in LUAD. As expected, the volume of the *TMPRSS2*-knockdown tumors had a significantly higher level of decreases than that of *TMPRSS2*-wildtype tumors after treatment with BMS-1, an inhibitor of PD-1/PD-L1 (Figure 5K); this result supports that knockdown of *TMPRSS2* can enhance the sensitivity of LUAD to the PD-1/PD-L1 inhibitor. Furthermore, the activities of CD8+ TILs and NK TILs markedly increased in *TMPRSS2*-knockdown tumors after treatment with BMS-1; they were significantly higher in *TMPRSS2*-knockdown than in *TMPRSS2*-wildtype tumors after treatment with BMS-1 (Figure 5L, M). These results support that the PD-1/PD-L1 inhibitor promotes immune elimination of tumor cells by inhibiting the exhaustion of CD8+ TILs and NK TILs in *TMPRSS2*-depleted LUAD.

To summarize, bioinformatics analysis revealed a negative correlation between TMPRSS2 abundance and immune infiltration levels in LUAD. Experimental results demonstrated that this relationship was a causal relationship. That is, reduced TMPRSS2 abundance can boost immune infiltration for LUAD.

## DISCUSSION

As a pivotal molecule in the regulation of SARS-CoV-2 invading human host cells, TMPRSS2 is attracting massive attention in the current SARS-CoV-2 pandemic [32-34]. Because SARS-CoV-2 has and is infecting large numbers of people, including many cancer patients, an investigation into the role of TMPRSS2 in cancer may provide valuable advice for treating cancer patients infected with SARS-CoV-2. Previous studies of TMPRSS2 in cancer mainly focused on its oncogenic role in prostate cancer [6-8]. In this study, we focused on LUAD, considering that it is the most common histological type in lung cancer and that the lungs are the primary organ SARS-CoV-2 attacks. In contrast to its oncogenic role in prostate cancer, TMPRSS2 plays a tumor suppressive role in LUAD, as we have provided abundant evidence. First, *TMPRSS2* downregulation correlates with elevated activities of many oncogenic pathways in LUAD, including cell cycle, mismatch repair, p53, and ECM signaling. Second, *TMPRSS2* downregulation correlates with increased tumor cell proliferation, stemness, genomic instability, and ITH in LUAD. Finally, *TMPRSS2* downregulation is associated with tumor advancement and worse survival in LUAD. Furthermore, both in vitro and in vivo experiments demonstrated that *TMPRSS2* downregulation markedly promoted the proliferation and invasion capacity of LUAD cells, supporting the tumor suppressor role of TMPRSS2 in LUAD.

Our bioinformatics analysis revealed significant negative associations between *TMPRSS2* expression and immune signatures, including both immune-stimulatory and immune-inhibitory signatures, in LUAD (Figure 1A). Nevertheless, *TMPRSS2* expression tended to have a stronger negative correlation with immune-inhibitory signatures than with immune-stimulatory signatures in LUAD (Figure 1B). The significant different levels of correlations of immune-stimulatory and immune-inhibitory signatures with *TMPRSS2* expression could be a factor responsible for the worse prognosis in LUAD patients with TMPRSS2 deficiency. Furthermore, the associations between TMPRSS2 and tumor immunity in LUAD were completely verified by both in vitro and in vivo experiments. That is, knockdown of *TMPRSS2* significantly increased tumor immunogenicity and immune cell infiltration in LUAD. On the other hand, both computational and experimental data showed that *TMPRSS2* downregulation significantly enhanced PD-L1 expression in LUAD. Because both inflamed tumor microenvironment and PD-L1 expression are determinants of cancer response to immunotherapy [35], *TMPRSS2*-depleted LUAD would respond better to immunotherapy than *TMPRSS2*-wildtype LUAD. This was supported by our in vivo experiments showing that *TMPRSS2*-knockdown tumors were more sensitive to the PD-1/PD-L1 inhibitor. Thus, *TMPRSS2* downregulation is a positive biomarker of immunotherapy for LUAD. In addition, because *TMPRSS2* downregulation often occurs in advanced LUAD, it indicates that advanced LUAD could benefit more from immunotherapy than early-stage LUAD.

TMPRSS2 inhibition has been indicated as a strategy for preventing and treating SARS-CoV-2 infection for the crucial role of TMPRSS2 in the SARS-CoV-2 invasion [33, 36]. However, our data suggest that this strategy may not be a good option for lung cancer patients in terms of the tumor suppressor role of TMPRSS2 in LUAD. Interestingly, we found that *TMPRSS2* displayed significantly higher expression levels in non-smoker than in smoker LUAD patients in four LUAD cohorts in which related data were available (Student’s *t* test, *p* < 0.05, FC > 1.5) (Figure 6A). This result indicates that non-smoker LUAD patients could be more susceptible to SARS-CoV-2 infection than smoker LUAD patients. As expected, non-smoker LUAD patients had significantly lower TMB and antitumor immunity than smoker LUAD patients (Figure 6B), consistent with findings from previous studies [37, 38].

**Figure 6.**
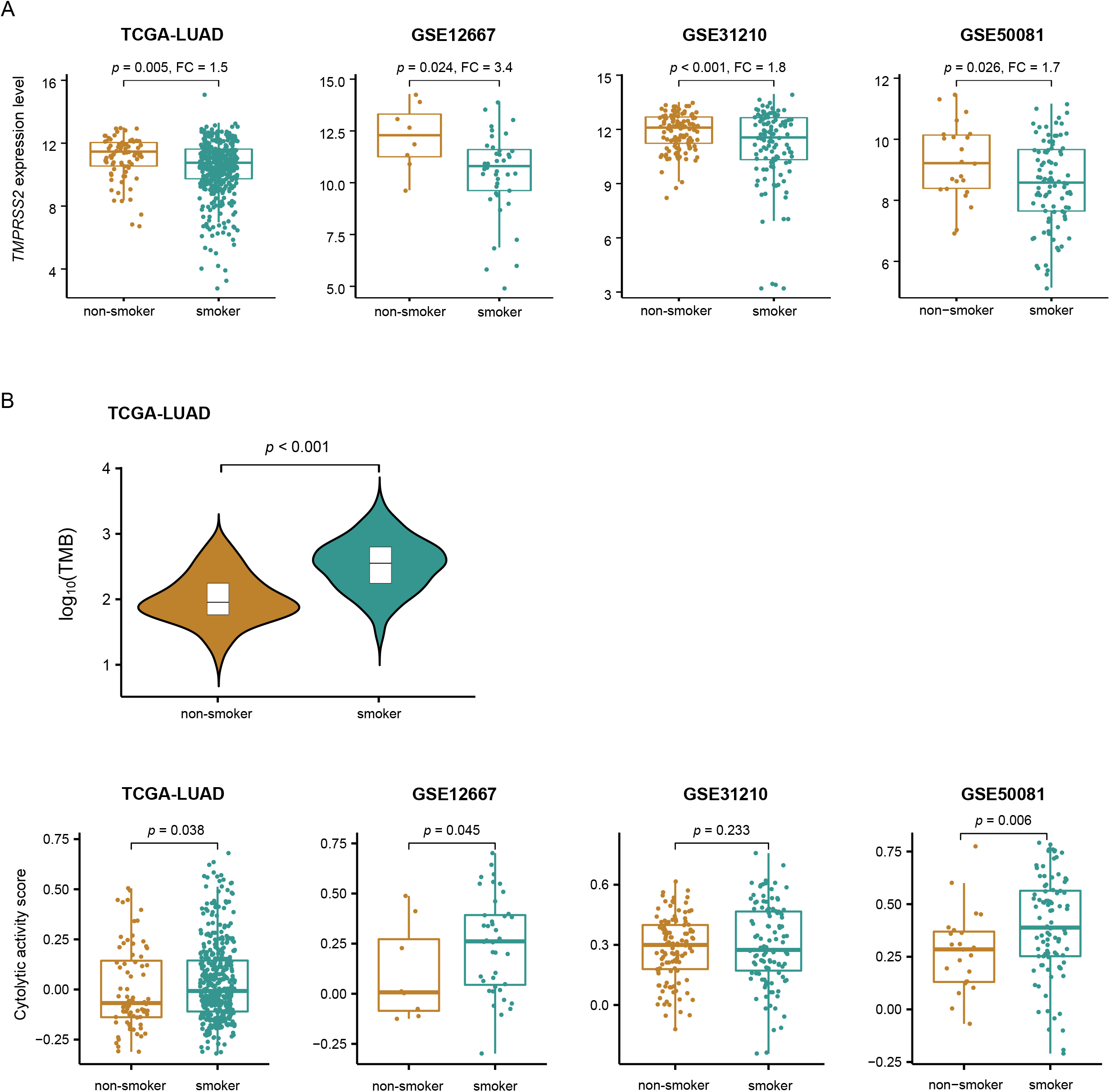
Comparisons of *TMPRSS2* expression levels, TMB, and immune signatures between non-smoker and smoker LUADs. Non-smoker LUAD patients showing significantly higher *TMPRSS2* expression levels **(A)** and lower TMB and immune signature scores **(B)** than smoker LUAD patients. The two-tailed Student’s *t* test and one-tailed Mann–Whitney *U* test *p* values are shown in **(A)** and **(B)**, respectively.

## CONCLUSIONS

TMPRSS2 is a tumor suppressor in LUAD, as evidenced by its downregulation correlated with increased genomic instability and ITH, tumor progression, and unfavorable clinical outcomes in LUAD. However, *TMPRSS2* downregulation is a positive biomarker of immunotherapy for LUAD. Our data provide implications in the connection between lung cancer and pneumonia caused by SARS-CoV-2 infection.

## Declarations

### Ethics approval and consent to participate

The study was done in accordance with both the Declaration of Helsinki and the International Conference on Harmonization Good Clinical Practice guidelines and was approved by the institutional review board.

### Consent for publication

Not applicable.

### Availability of data and material

The five LUAD genomic datasets were obtained from the Genomic Data Commons Data Portal (https://portal.gdc.cancer.gov/) and the Gene Expression Omnibus (https://www.ncbi.nlm.nih.gov/geo/).

### Authors’ contributions

**Zhixian Liu**: Validation, Formal analysis, Resources, Investigation, Data curation, Visualization, Writing -original draft, Funding acquisition. **Zhilan Zhang**: Software, Formal analysis, Investigation, Data curation, Visualization. **Qiushi Feng**: Software, Formal analysis, Visualization. **Xiaosheng Wang:** Conceptualization, Methodology, Resources, Investigation, Writing - original draft, Writing - review & editing, Supervision, Project administration, Funding acquisition.

## List of Abbreviations

ACE2: angiotensin-converting enzyme 2
CCK-8: the Cell Counting Kit-8
DAPI: 4’,6- diamidino-2-phenylindole
DDR: DNA damage repair
DFS: disease-free survival
ECM: extracellular matrix
FC: fold change
FDR: false discovery rate
GO: gene ontology
GSEA: gene set enrichment analysis
HRD: Homologous recombination deficiency
ICIs: immune checkpoint inhibitors
ITH: intratumor heterogeneity
LUAD: lung adenocarcinoma
LUSC: lung squamous cell carcinoma
MDSCs: myeloid-derived suppressor cells
OS: overall survival
PI: proximal-inflammatory
PP: proximal-proliferative
RT-PCR: Real-Time PCR
S: spike glycoprotein
SARS-CoV-2: severe acute respiratory syndrome coronavirus 2
siRNA: small interfering RNA
ssGSEA: single-sample gene-set enrichment analysis
TCGA: The Cancer Genome Atlas
TILs: tumor-infiltrating lymphocytes
TMB: tumor mutation burden
TMPRSS2: transmembrane protease serine 2
TRU: terminal respiratory unit
WGCNA: weighted gene co-expression network analysis

## Supplementary data

**Table S1**. A summary of the datasets analyzed.

**Table S2**. The marker gene sets of immune signatures, pathways, and tumor phenotypes.

**Table S3**. The genes with strong positive and negative expression correlations with *TMPRSS2* in the TCGA-LUAD cohort.

